# Selection and adaptive introgression guided the complex evolutionary history of the European common bean

**DOI:** 10.1101/2022.09.28.509856

**Authors:** Elisa Bellucci, Andrea Benazzo, Chunming Xu, Elena Bitocchi, Monica Rodriguez, Saleh Alseekh, Valerio Di Vittori, Tania Gioia, Kerstin Neumann, Gaia Cortinovis, Giulia Frascarelli, Ester Murube, Emiliano Trucchi, Laura Nanni, Andrea Ariani, Giuseppina Logozzo, Jin Hee Shin, Chaochih Liu, Liang Jiang, Juan José Ferreira, Ana Campa, Giovanna Attene, Peter Laurent Morrell, Giorgio Bertorelle, Andreas Graner, Paul Gepts, Alisdair Robert Fernie, Scott Allen Jackson, Roberto Papa

## Abstract

Domesticated crops have been disseminated by humans over vast geographic areas. After 1492, the common bean (*Phaseolus vulgaris* L.) was introduced in Europe. Here, we combine whole-genome profiling, metabolic fingerprinting and phenotypic characterisation, and we show that the first common bean cultigens successfully introduced into Europe were of Andean origin, after Francisco Pizarro’s expedition to northern Peru in 1529. We show that hybridisation, selection and recombination have shaped the genomic diversity of the European common bean in parallel with political constraints. There is clear evidence of adaptive introgression into the Mesoamerican-derived European genotypes, with 44 Andean introgressed genomic segments shared by more than 90% of European accessions and distributed across all chromosomes except PvChr11. Genomic scans for signatures of selection highlight the role of genes relevant to flowering and environmental adaptation, suggesting that introgression has been crucial for the dissemination of this tropical crop to the temperate regions of Europe.

## Introduction

Following the process of domestication, crops were spread by humans over vast geographic areas, where they adapted to new and often extreme environments^1^. The Columbian Exchange^2^ started in 1492 with the transatlantic journey of Christopher Columbus. This large-scale set of reciprocal biological introductions between continents provides a paradigm for the rapid adaptation of crop plants to changing environments. Changes in flowering time and photoperiod sensitivity were selected in parallel in the common bean (*Phaseolus vulgaris*), maize (*Zea mays*), potato (*Solanum tuberosum*), to name a few crops that have undergone selection for these adaptive traits^1, 3^. Among crops originating from the Americas, the common bean was rapidly adopted and successfully disseminated across Europe^4^ and it is now possible to identify local European varieties with Andean and Mesoamerican origins^5–10^. Nowadays, common bean, as other food legumes, is crucial for main societal challenges and to promote the transition to plant-based diets^11^.

The introduction of the common bean to Europe from two distinct centres of origin offered an opportunity for widespread genepools hybridisation and recombination^9^. Studies of common bean evolution in Europe can exploit the parallel domestication processes and the major genetic differences between the two American genepools. This provides an ideal model to study the role of introgression during the adaptation of common bean accessions to European environments^12^.

In this work, we combine whole-genome analysis and metabolic fingerprinting in 218 common bean landraces, integrated with genome-wide association (GWA) to characterise the genetic basis of multiple traits, including flowering time and growth habit in different environments with contrasting photoperiod conditions. We have used the combined results to characterise the effects of selection and inter-genepool introgression, and to test the occurrence of adaptive introgression associated with the development and adaptation of common bean accessions in Europe.

## Results

### The common bean population structure reveals pervasive admixture in Europe

Here, we characterized common bean landraces (*P. vulgaris*) (Supplementary Data 1), performed multi-location field experiments and trials under greenhouse-controlled conditions, and carried out classical and molecular phenotyping (i.e., metabolomics) (see Supplementary Notes 1-6, Supplementary Fig. 1-4 and Supplementary Tab. 1-4). We performed whole-genome sequencing (WGS) and a detailed summary on sequencing results is provided in Supplementary Notes 7-11 (see also Supplementary Fig. 5-10, Supplementary Tab. 5-6 and Supplementary Data 2-3).Using ADMIXTURE^13^ (Supplementary Notes 12-15), we reconstructed the ancestry of 218 single-seed-descent (SSD) purified accessions from the Americas (104 accessions including 66 pure American accessions showing low admixture between gene pools, qi > 99%) and from Europe (n=114), based on nuclear (Supplementary Fig. 11-19) and chloroplast (Supplementary Note 16 and Supplementary Fig. 20-22) genetic variants (Fig. 1a–d). The accessions were spatially interpolated to investigate their geographic distribution in Europe (Supplementary Note 17, Fig. 1e, and Supplementary Fig. 23) and we proposed a model for their introduction in the Old Continent (Fig. 1f). For common beans originating from the Americas, subdivision into the highly differentiated Andean and Mesoamerican populations was consistent with previous studies^12, 14, 15^ (Fig. 1a). Next, we followed the nested procedure previously used by Rossi et al.^16^ to investigate the population structure within each American gene pool (Supplementary Note 15). Again, using ADMIXTURE^13^, we identified two main Mesoamerican groups (M1 and M2) and three main Andean groups (A1, A2 and A3) (Fig. 1b, c). In the centres of domestication, there was little evidence of admixture between genepools.

**Figure 1.**
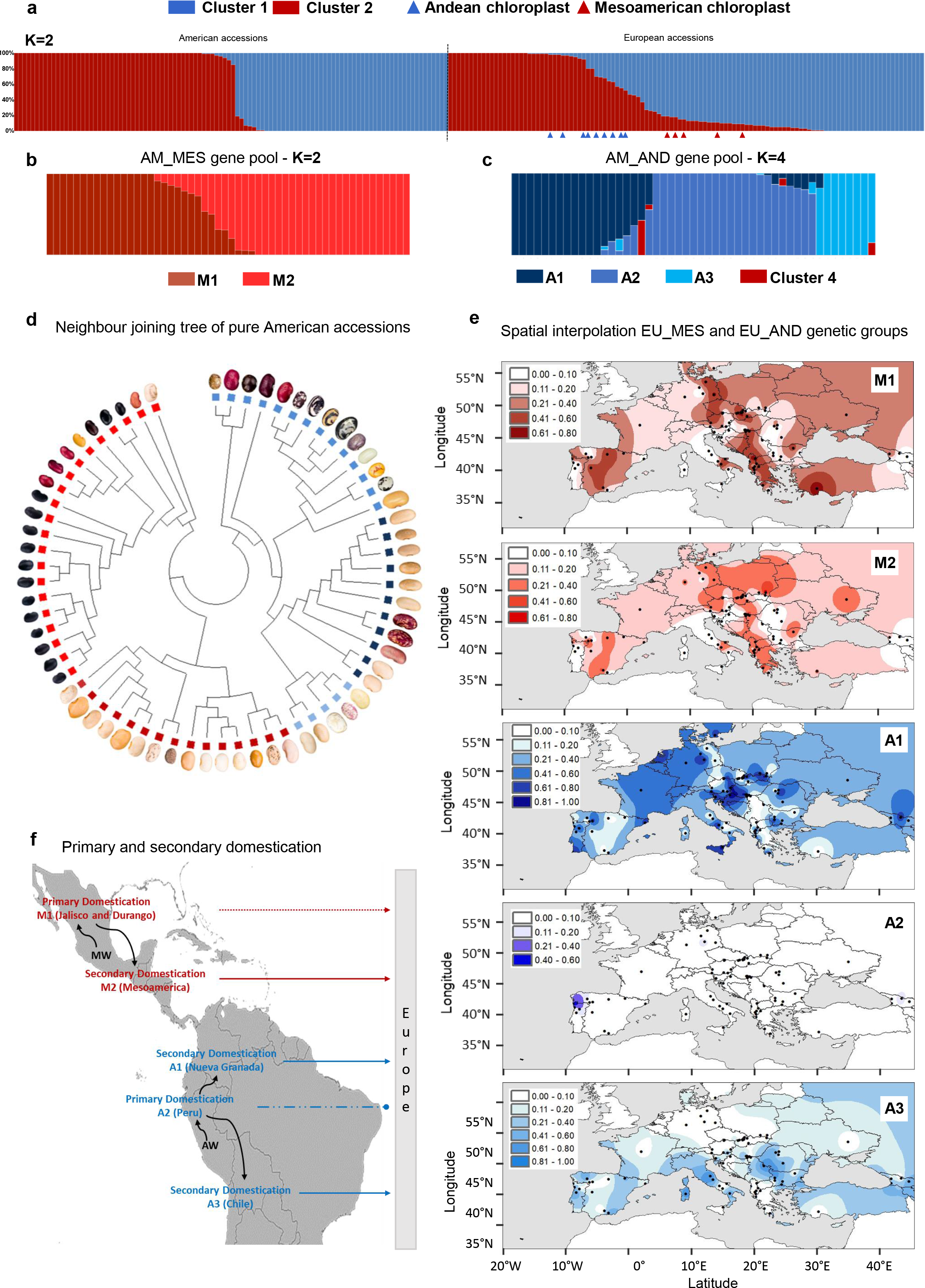
Population structure of common bean in America and Europe. **a,** Admixture analysis (K = 2) showing inferred ancestry in the American (AM; left) and European (EU; right) accessions, with the identification of two gene pools (identified as clusters 1 and 2) that show correspondence to the two main common bean gene pools based on our passport data (cluster 1 – Andean, cluster 2 – Mesoamerican), and several intermediates and admixed genotypes in Europe. The chloroplast ancestry assignment is shown for the accessions (triangles below the chart) when not consistent with the nuclear assignment **b,** Admixture plots for the AM Mesoamerican accessions (K = 2) grouped by geographic origin (i.e., latitude and state), which identifies two main subgroups (M1 and M2). **c,** Admixture plots for the AM Andean accessions (K = 4) grouped by geographic origin (i.e., latitude and state), which identifies three Andean genetic subgroups (A1, A2 and A3). A fourth cluster in four accessions, based on the whole-set ADMIXTURE analysis (K = 2), was induced by the occurrence of Mesoamerican alleles with AM_M1/AM_M2 components (see also supplementary Note 15). **d,** Neighbour-joining tree and seed pictures of the 66 pure American accessions. **e,** Spatial interpolation of the geographic distributions of the EU Mesoamerican (M1 and M2) and EU Andean (A1, A2 and A3) ancestry components in Europe, as inferred by ChromoPainter analysis. Maps were designed using the map tools implemented in different R packages, such as *spatial*, *maps*, *fields*, *maptools*, *raster*, *rgdal*. **f,** Primary and secondary domestications of Mesoamerican and Andean genetic groups/races in America. Loss of photoperiod sensitivity during secondary domestication was a relevant factor for the introduction of the Andean A1/Nueva Granada and A3/Chile and for the Mesoamerican M2/ Mesoamerica ancestries in Europe (solid arrow). Genetic group M1 (Durango-Jalisco race) was successfully introduced into Europe after introgression from other genetic groups characterised by little or no photoperiod sensitivity (dashed arrow). Genetic group A2 (Peru race) was not introduced into Europe due to its high photoperiod sensitivity (discontinuous and truncated line). Source data (a,b,c) are provided as a Source Data file.

Among the European accessions, nuclear variants allowed us to identify several admixed genotypes (35 EU accessions with more than 10% and 18 with more than 20% of the genome attributed to the other gene pool based on the admixture) (Fig. 1a). In a few European accessions (n = 14), the nuclear and chloroplast assignments were inconsistent (Fig. 1a): considering only accessions with introgressed genome >70% (n=11), the Andean chloroplast genome was combined with a Mesoamerican nuclear genome (n =6) or *vice versa* (n=5). This suggests, at least in some cases, the occurrence of chloroplast capture^17^ as a result of inter-genepool hybridisation and subsequent backcrossing. Moreover, when we explored the molecular phenotypic diversity of the American and European accessions (Supplementary Note 5), the metabolomic fingerprint (the molecular phenotypic space expressed as principal component 1 from 1493 putative secondary metabolites with a high hereditability of H^2^ > 0.65) confirmed the admixture scenario. Several intermediate phenotypes between Mesoamerican and Andean accessions were observed in European landraces, but these were absent in accessions collected in the Americas (Fig. 2a-c). Notably, there was a significant correlation between the admixture coefficients and principal component 1 for both the American and the European accessions, indicating a tight relationship between the phenotypic and genotypic differences due to the genepool structure. This included a reduced difference in Europe due to admixture, particularly in the accessions of Mesoamerican origin. The occurrence of pervasive admixture among gene pools characterises also other centres of cultivation of common bean (See Supplementary Note 17 and Supplementary Fig. 24), and it is supported in Europe also by our phylogenetic network analysis (See Supplementary Note 18 and Supplementary Fig. 25-26 for more details).

**Figure 2.**
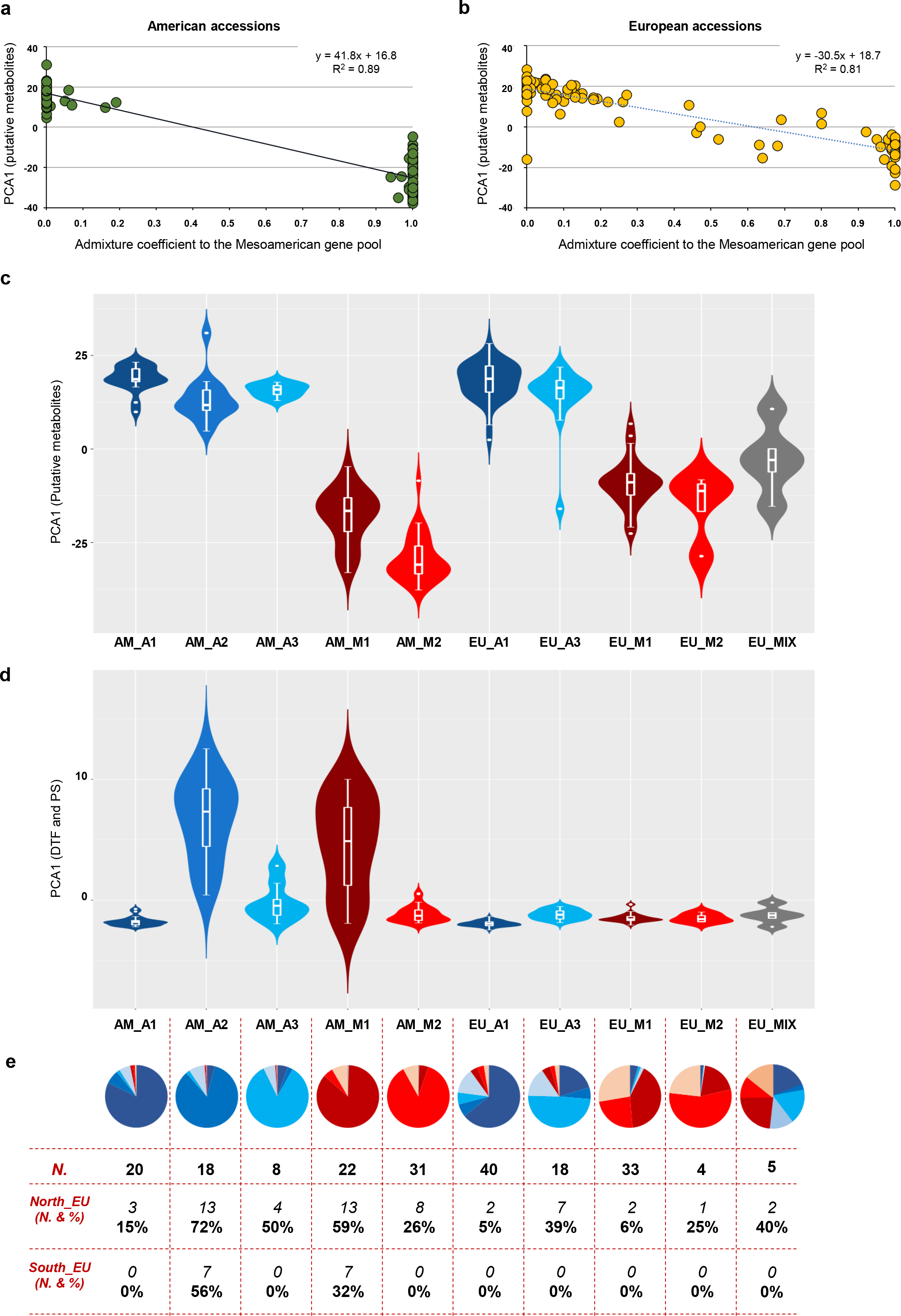
Genetic structure, molecular phenotyping and flowering data. **a, b,** Molecular phenotypes (PCA1 from 1493 putative secondary metabolites, H^2^ > 0.65 over the entire dataset) of (a) 94 American accessions and (b) 96 European accessions confirm the subdivision into the two main groups based on the admixture coefficient (derived from nuclear genomic data, K = 2). Intermediate phenotypes and genotypes are seen in Europe. **c,** Violin plots showing the distribution of PCA1 values related to secondary metabolites showing high heritability (H^2^ > 0.65) by genetic subgroups in the American and European accessions. PCA1 was used as a representative molecular phenotype, and it explains 25.7% of the total variance for these traits. N. biologically independent samples, AM_A1 (19), AM_A2 (15), AM_A3 (8), AM_M1 (20), AM_M2 (31), EU_A1 (39), EU_A3 (17), EU_M1 (31), EU_M2 (4), EU_MIX (5). **d,** Violin plots showing the distribution of PCA1 values related to the days to flowering (DTF) and photoperiod sensitivity (PS) by genetic subgroups in the American and European accessions. PCA1 was used as a representative phenotypic trait for DTF and PS, and it explains 68.8% of the total variance for these traits. N. biologically independent samples, AM_A1 (20), AM_A2 (18), AM_A3 (8), AM_M1 (22), AM_M2 (31), EU_A1 (40), EU_A3 (18), EU_M1 (33), EU_M2 (4), EU_MIX (5). **e,** Proportions of the genetic memberships – P(AM_A1), P(AM_A2), P(AM_A3), P(AM_M1), P(AM_M2), P(SAND), and P(SMES) – inferred from the donor accessions and composing the American and European accessions (grouped as mainly AM_A1, AM_A2, AM_A3_AM_M1, AM_M2, EU_A1, EU_A3, EU_M1, EU_M2, and EU_MIX) are shown in the pie charts below the corresponding groups and flowering data (number and percentage of individuals with delayed or no flowering) in northern and southern Europe, related to the corresponding groups. c,d, box plots represent minimum, first quartile, median, third quartile and maximum. Source data (c,d) are provided as a Source Data file.

Finally, to provide insight into the population structure of the American pure accessions, we considered their passport data (geographic distribution and country of origin) and phenotypic data on growth habits and photoperiod sensitivities. We identified a clear correspondence between the genetic groups in our sample from the Americas and the well-known common bean eco-geographic races^18^: M1 corresponded to the higher-altitude Durango and Jalisco races, which originated primarily in northern and southern Mexico, respectively; M2 corresponded to the lower-altitude Mesoamerican race, which is mostly photoperiod-insensitive and is distributed in lowland Mexico, in Central America and in the Caribbeans ; A1 corresponded to the generally photoperiod-insensitive Nueva Granada race; A2 corresponded to the Peru race, which includes entries with vigorous climbing growth habits and photoperiod sensitivity; and A3 corresponded to the Chile race, which has also been identified in archaeological samples from northern Argentina that date from 2500 to 600 years ago^19^. The identification of these well-defined ancestral genetic groups in the Americas offers a robust basis to study the inter-genepool and inter-race introgression that may have facilitated adaptation to European environments.

### Asymmetric introgression and recombination between genepools underlie European common bean adaptation

Given the presence of admixed individuals, we studied the inter-genepool hybridisation and introgression pattern associated with the evolutionary history of the common bean in Europe by genetic assignment at the chromosome level in ChromoPainter v2.0^20^ (Supplementary Note 19). When compared to admixture analysis, this can provide information on recombination between markers and the size of regions that can be attributed to different ancestries. The 66 pure American accessions (Supplementary Data 4), from the five genetic groups identified using admixture, due to their low levels of admixture, were used as donor (reference/founder) populations for the chromosome painting of the European genotypes. On this basis, and for each European genotype, we attributed all the single-nucleotide polymorphisms (SNPs) and chromosomal regions to particular ancestries, taking into account within-accession recombination breakpoints. Using this approach, we were also able to detect recombination events between genepools at the whole-genome level, even in accessions that showed < 1% introgression. Overall, 71 European accessions were attributed to the Andean gene pool (EU_AND) and 43 were assigned to the Mesoamerican gene pool (EU_MES), in agreement with the admixture analysis (Pearson correlation: r = 0.99, p < 0.01; Supplementary Fig. 27) and confirming previous knowledge about the prevalence of Andean genotypes in Europe^5, 21^. Globally, the inferred amount of per-accession introgressed material differed between the EU_MES and EU_AND accessions (two-sided K-S test, p = 3.3×10^−3^), showing median proportions of 4.7% and 9.2%, respectively. In the EU_MES accessions, 0.01–44.9% of the genome had introgressed from the other genepool, with only one EU_MES accession showing < 1% genome introgression (Fig. 3a; see also Supplementary Note 19). These proportions were similar in the EU_AND samples, ranging from 0% to 42.2% (Fig. 3a), although two EU_AND accessions showed no introgression and 10 EU_AND accessions showed < 1% introgression from the other genepool (Fig. 3a). The pervasive effect of admixture in European individuals was confirmed by the presence of several accessions with > 20% of their genome acquired by introgression in both the EU_MES (8 of 43 accessions, 18.6%) and EU_AND (11 of 71 accessions, 15.5%) groups (Fig. 3a). The inferred ancestry per each chromosome is represented in Supplementary Fig. 28-38.

**Figure 3.**
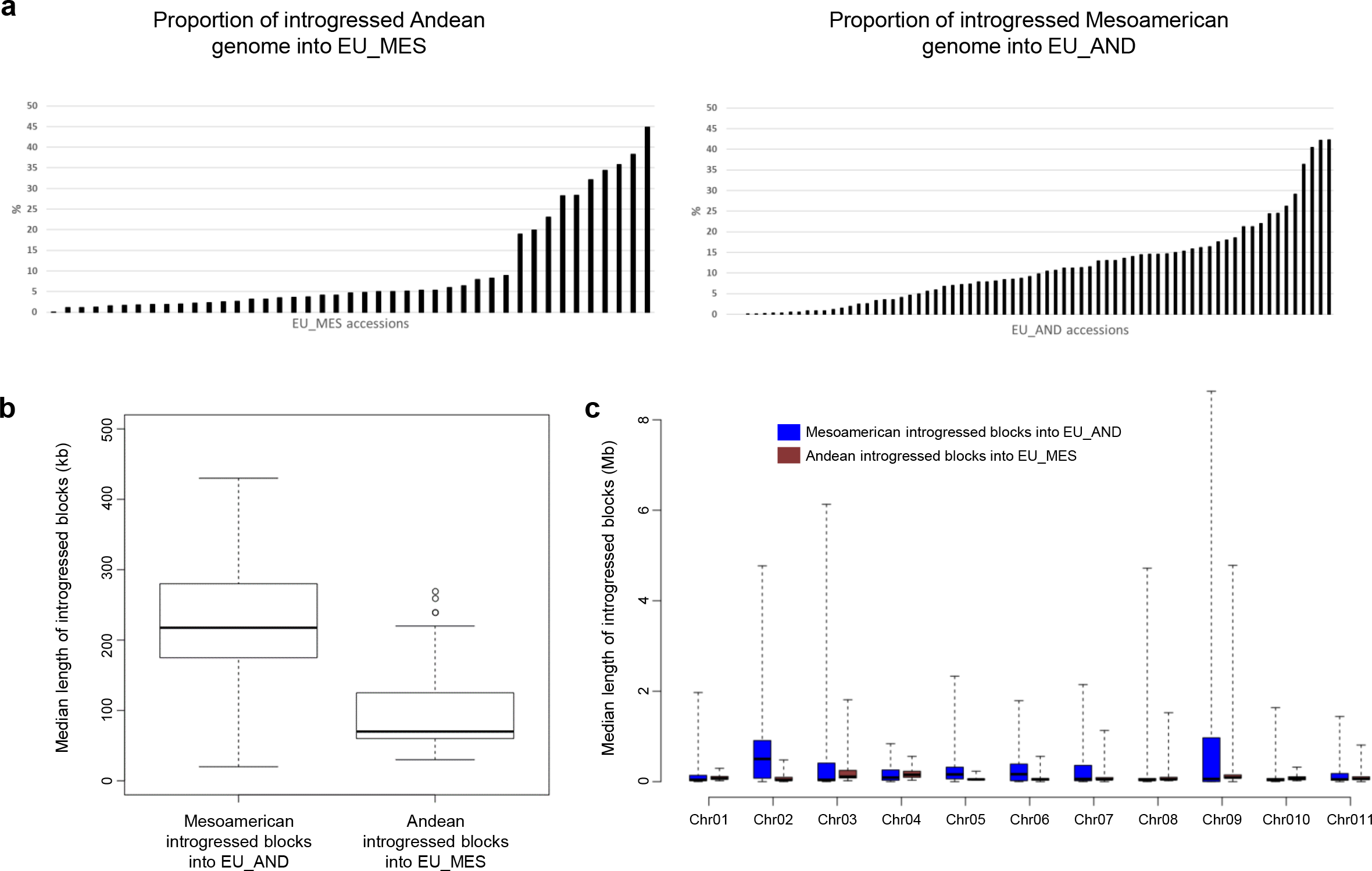
Mapping introgression in the European common bean using ChromoPainter. **a,** Proportion of introgressed genome in the Mesoamerican (EU_MES; n = 43) and Andean (EU_AND; n = 71) groups. **b, c,** Boxplots showing the median length of the introgressed blocks identified in each of the EU_AND and EU_MES accessions across all of the chromosomes (b) and the median length of the introgressed blocks identified in each of the EU_AND and EU_MES individuals by chromosome (c). b,c, box plots represent minimum, first quartile, median, third quartile and maximum. Sample size (N. accessions), (b,c; EU_AND=71, EU_MES=43). Source data (a,b) are provided as a Source Data file.

The median length of the introgressed genomic segments was higher for the EU_AND accessions (EU_AND = 217 kb, EU_MES = 70 kb; Mann-Whitney test, P=7.22×10^-10^, Fig. 3b,c), with more extended regions introgressed into EU_AND particularly on chromosomes PvChr02, PvChr05, PvChr06 and PvChr09 (Fig. 3b,c). We obtained very similar results when we repeated the analysis by excluding six accessions showing an admixture proportion >=40% (Supplementary Fig. 39). The EU_AND accessions carried longer Mesoamerican introgressed haplotypes; the maximum length of the introgressed genomic blocks measured in the EU_AND chromosomes was always higher than what present in EU_MES individuals (Supplementary Tab. 7), supporting the more recent introgression of Mesoamerican genome fragments into the Andean genotypes compared to the opposite direction. When we estimated the timing of the introgression among the two gene pools in Europe, we confirm the more recent introgression from the Mesoamerican to the Andean gene pool, while the Mesoamerican gene pool was introgressed earlier (Supplementary Note 20, Supplementary Fig. 40). This estimated introgression time is compatible with historical data and with an earlier successful introduction of the Andean gene pool in Europe. Several genomic regions that carry haplotypes with a specific Andean ancestry are near fixation in the European accessions. Here, when seeking regions that may have been subject to natural selection, playing a role in the adaptation, we identified regions putatively under selection in Europe (Supplementary Note 21, Supplementary Tab. 8 and Supplementary Data 5). Signatures of selection were detected among the regions with Andean ancestry nearly fixed in the European accessions (e.g., position 46 Mb on chromosome Pv01, which carries the *OTU5* locus, that may be involved in the phosphate starvation response; Supplementary Note 19, Supplementary Fig. 28; and on position 37.9 Mb on chromosome Pv09 which carries the *LHY* locus)).

Our results indicate that the first cultigen successfully disseminated across Europe was composed of Andean types. This is shown by the smaller introgression segments of Andean origin and the higher frequencies of Andean-derived common bean accessions in Europe. Our data are also consistent with available historical records. Indeed, the first unambiguous evidence for the introduction of common bean in Europe points to Andean cultivars^22^ probably introduced into Spain by Francisco Pizarro in 1529 following the exploration of Peru. Piero Valeriano Bolzanio received common bean seeds from Giuliano de Medici (Pope Clement VII, 1523–1534), which had been donated to the same pope by Emperor Charles V’s Spanish emissaries from Sicily (where the bean seeds were harvested). The very detailed writing of Piero Valeriano Bolzanio refers to common bean seeds, describing in depth several phenotypic traits supporting their Andean origin, as also recently proposed^23^. Valeriano documented his efforts, along with a network of collaborators in the north-east of Italy, Slovenia and Dalmatia, to grow and reproduce beans starting in 1532^22^, with the first report of a putative Mesoamerican genotype in Europe dated 1542^23^. Historical information and timelines, together with our data showing asymmetric introgression, suggest an earlier successful introduction and spread of the Andean genepool into Europe. This may also explain the high frequency of A1/Nueva Granada Andean ancestries (Fig. 1e) in Sicily, the south and north-east of Italy, Slovenia and Croatia, because they could have been among the first European areas to cultivate common bean with the earliest introduced Andean genotypes probably from the Nueva Granada race.

Adaptive differences among common beans in the New World may also have influenced the distributions in Europe. For example, M1/Durango-Jalisco genotypes can be particularly photoperiod sensitive, and may therefore have failed to adapt well to many European environments, thus, limiting their dissemination, particularly in central and northern Europe (Fig. 1e). In contrast, southern Spain, southern Italy, Sicily, North Africa, Madeira Island, and the Canary Islands are characterised by mild winters. In these environments, photoperiod-sensitive and late-flowering genotypes, or those adapted to warmer conditions, may have easily completed the crop cycle. As also reported by others^5–24^, we found that Mesoamerican genotypes are more frequent in specific European regions, particularly in south-eastern Europe (Fig. 1e), which also suggests that the history of their introduction may have contributed to their current distribution. As with the role of Charles V and Pope Clement VII in the early dissemination of the Andean beans, the political subdivision of Europe and the Mediterranean basin in the 16^th^ century may have influenced the dissemination of the Mesoamerican genepool. The Ottoman Empire dominated the southern shores of the Mediterranean, the Nile Basin, the Red Sea into eastern Africa, and south-eastern Europe, spanning the area from modern-day Greece to Austria. The prevalence of Mesoamerican genotypes in eastern Africa and China^24, 25^ may reflect their initial introduction into Africa from Spain during the Ottoman Empire, which extended its rule in north-eastern Africa and controlled the exchange of goods with China through the Silk Road. Although additional comparative studies between European and Chinese centres are required, our hypothesis is compatible with our results from a *de-novo* admixture analysis applied to Chinese landraces^25^, reported in Supplementary Note 17 (Supplementary Fig. 24). The importance of political/cultural factors associated with the dissemination of common bean genotypes in Europe is compatible with the lack of significant spatial and ecological patterns between genetic, geographic and ecological distances. Indeed, the routes of dissemination based on cultural and political factors are often independent of geographic and environmental distances, making the occurrence of correlations between genetic distances and geographical or environmental differences less likely^26–30^ (see also Supplementary Note 22).

### Analysis of the Environmental associations

We used the geographic distribution of the five ancestral components inferred by ChromoPainter in an association analysis with biogeographical variables (Supplementary Note 22 and Supplementary Data 6). Ancestral components of A3/Chile negatively correlated with latitude (Supplementary Tab. 9; r = – 0.35, p = 0.0001) and were never observed above the 47^th^ parallel (Fig. 1e). Moreover, A3/Chile component was associated with warmer climates, particularly the maximum temperature in September (Supplementary Tab. 9; r = 0.29, p < 0.002). Although A3/Chile did not appear any more photoperiod-sensitive than A1/Nueva Granada (see Supplementary Note 23 on flowering and metabolomics variability among genetic groups; Supplementary Tab. 10-11), some American A3/Chile individuals tend to flower later (Fig. 2d and Supplementary Fig. 41) at higher latitudes when grown in Europe (Fig. 2e). Here, we suggest that although A3/Chile was successfully introduced in Europe, a residual sensitivity to the photoperiod might be still preserved in some European genotypes mainly belonging to this ancestry, that show delayed flowering at certain latitudes (Fig 2e), that may have also influenced the dissemination of this common bean ancestry in Europe (Fig. 1e,f). However, compared to the Mesoamerican genotypes, A3/Chile was more uniformly distributed in Europe across different longitudes (Fig. 1e), which also supports the earlier introduction of Andean genotypes. Only a few weak associations with environmental variables were detected for the other genetic groups (Supplementary Note 22).

### Analysis of genetic diversity in the European common bean

To disentangle how inter-genepool hybridisation have shaped the genetic diversity of the European common bean, and given the evidence of widespread admixture in Europe, we developed a masked dataset of European accessions by filtering out all introgressed alleles or those with an ambiguous assignment. Detailed results on the analysis of genetic diversity are reported in Supplementary Notes 24-26 (see also Supplementary Fig. 42-59 and Supplementary Tab. 12-13).The development of a masked dataset allowed us to consider nucleotide diversity using the frequencies of two reconstructed non-admixed populations of Andean and Mesoamerican origin. From each European genotype, all the Andean SNPs were separated from the Mesoamerican SNPs and included in the two masked datasets. Based on the unmasked and masked datasets, American common bean accessions showed moderately higher nucleotide diversity than European accessions, apparently due to the introduction bottleneck in Europe (Fig. 4a,b). Moreover, when compared to the Mesoamerican gene pool, the Andean gene pool showed an overall lower diversity in the primary centres of domestication (Americas) using both the masked and unmasked datasets (Fig 4a, b). This confirms that the diversity of the Andean germplasm at the centre of origin might still reflect the bottleneck that occurred in the Andean wild populations during the expansion into South America before domestication^31^, as reflected in the domesticated pool^32^. Indeed, we detected ∼70% lower diversity (θ_π_/bp) in the Andean compared to the Mesoamerican accessions. Very similar results were obtained when we repeated the analysis by excluding six accessions showing an admixture proportion >=40% (Supplementary Note 25).

**Figure 4.**
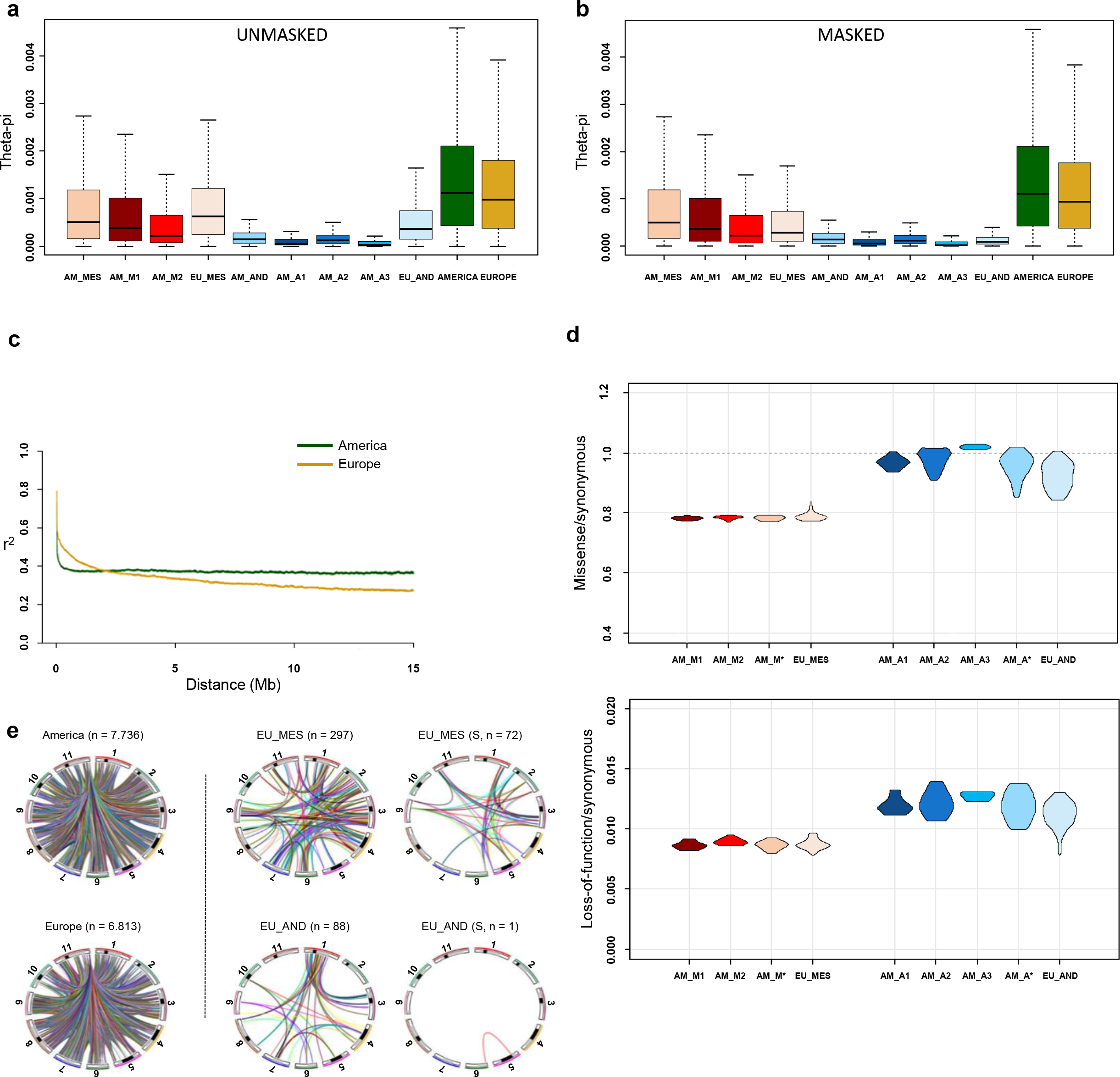
Boxplots of θπ averaged over 100-kb non-overlapping sliding windows, linkage disequilibrium (LD) decay and inter-chromosomal linkage disequilibrium and genetic load. **a,** Genetic diversity computed using whole chromosomes and the unmasked dataset. **b,** Genetic diversity computed after the admixture masking process using whole chromosomes and LD decay according to the physical distance. **c,** Comparative LD decay in the American and European accessions. **d,** Genome-wide measure of genetic load in the American and European accessions. The ratios are shown for missense (up) and loss-of-function (down) over synonymous mutations in the different groups. AM_M* and AM_A* are the admixed American accessions (not pure American individuals). **e,** Private inter-chromosomal LD in American and European accessions (left), in the Mesoamerican and Andean European accessions (middle), and considering genomic regions under selection (S) in the Mesoamerican and Andean European accessions (right) a,b, box plots represent minimum, first quartile, median, third quartile and maximum. a,b, N=5134 genomic windows. Source data (a,b,d) are provided as a Source Data file.

When American and European genetic diversities were compared within each genepool using the unmasked dataset (AM_AND *vs* EU_AND and AM_MES *vs* EU_MES), due to the admixture, European diversity was always higher than American diversity, but the opposite was found when using the masked dataset (Fig. 4b). In other words, we show how the Andean common beans from Europe are more diverse than those from America because of admixed ancestry with the Mesoamerican gene pool, as seen by comparing the genetic diversities of the unmasked and masked datasets (EU_AND). This comparison of the estimated levels of genetic diversity in Europe reflects the key role of inter-genepool hybridisation and recombination in shaping the diversity of the European common bean. The genetic diversity was higher in M1/Durango-Jalisco than M2 Mesoamerica accessions in Americas, and also in A2/Peru than A1/Nueva Granada accessions in Americas, whereas the amount of diversity in A3/Chile accessions was very low. Combined with the neighbourhood-joining tree shown in Fig. 1d, this indicates that the A2/Peru and M1/Durango-Jalisco races have been probably the earlier domesticated Andean and Mesoamerican populations, from which the other races arose by secondary domestication associated with the loss of photoperiod sensitivity (Fig. 1f). Indeed, earliness and loss/reduction of photoperiod sensitivity were important traits under selection during the expansion of the common bean in Europe. This is also suggested by our test for the occurrence of selection for flowering during the introduction of common bean in Europe (see also Supplementary Note 23). The genetic differentiation between American and European accessions for flowering time (PC1 based on flowering data across different European field and greenhouse trials) was measured using *Q* ^33^. The *Q_ST_* for flowering was compared with the distribution of the *Q_ST_* for highly hereditable metabolites and with the *F_ST_* distribution of the SNPs. We show that the *Q_ST_* for the flowering is in the top 97.5% of distribution of the *Q_ST_* for highly inheritable metabolites, being also an outlier (99.5%) compared to our *F_ST_* distribution, suggesting that flowering is likely a candidate trait that underwent a selection process^33, 34^. Considering the Andean genepool, the successful introduction in Europe was connected to the domestication pattern at the centre of origin. The earlier, photoperiod-sensitive domesticated genotypes were less successfully disseminated in Europe. Indeed, the relationship between the American and European genetic groups of Andean origin (as defined by ChromoPainter), coupled with the phenotypic data for flowering (Fig. 2d), shows that the A2/Peru race was more photoperiod-sensitive and was not introduced into Europe successfully, due to the lack of adaptation (other than a single highly admixed accession, qA2 = 43.6%). In contrast, the remaining Andean genetic groups (A1/Nueva Granada and partially A3/Chile) became widespread in Europe. A different scenario was seen for the Mesoamerican genotypes, especially M1/Durango-Jalisco, where introgression appears to have been an important element in the dissemination of the common bean in Europe (Fig. 1f, 2d,e). M1/Durango-Jalisco showed very high levels of admixture in the European material due to introgression from the M2/Mesoamerica and the A1/Nueva Granada and A3/Chile (Fig. 2e), which likely contributed to reduced photoperiod sensitivity compared to the American Durango-Jalisco counterpart (AM_M1) (Fig. 2d), supporting its dissemination throughout Europe (Fig. 1e,f).

For the Andean genotypes, both the diversity pattern and photoperiod sensitivity (Fig. 2d) suggest at least two domestication steps occurred: primary domestication of photoperiod-sensitive populations (A2/Peru) and secondary domestication characterised by reduced photoperiod sensitivity (A3/Chile and particularly A1/Nueva Granada). This indicates that secondary domestication^35^ was necessary for the successful dissemination of the Andean common bean in Europe (Fig 1f). For the Mesoamerican genotypes, an open question is where and when the introgression from the Andean genepool occurred. We suggest this is likely to have happened in southern Europe and along the southern Mediterranean shore, where the warmer climate in winter may have favoured the Mesoamerican genotypes.

The average linkage disequilibrium (LD) decay in accessions from Europe and the Americas (Fig. 4c) is consistent with the historical differences between the genepools and the effects of high inter-genepool hybridisation and introgression at the whole-genome scale in Europe (see also Supplementary Note 27). Admixture in Europe increased the molecular diversity (i.e., effective population size). It also generated new genome-wide admixture LD due to new combinations of alternative alleles in each genepool. Accordingly, inter-genepool hybridisation followed by recombination reduced LD at a long distance but, as expected, had a limited effect on LD decay at short distances because regions are directly inherited from the source populations^36^. When we compared the American and European accessions, LD decay was much faster over short distances (< 1.5–2.0 Mb) in American genotypes. In contrast, there was faster LD decay over greater distances (> 3 Mb) in European populations (Fig. 4c). This reflects higher historical rates of recombination in the American genotypes over short distances and the effect of recombination due to inter-genepool introgression in Europe over long distances. A similar pattern was seen when the Mesoamerican and Andean genepools were analysed separately (Supplementary Fig. 60). However, the Andean accessions were characterised by higher baseline LD levels. Indeed, the AM_MES and AM_AND populations reached *r*^2^ = 0.2 at ∼500 kb and ∼1 Mb, respectively, whereas *r*^2^ = 0.2 was reached at ∼1.1 and ∼3.5 Mb for the EU_MES and EU_AND samples, respectively.

### Synonymous and missense mutations

The ratios between missense and loss-of-function mutations over synonymous mutations were calculated to reveal patterns of genetic load across genepools and continents (see Supplementary Note 28). The total amount of synonymous, missense, stop-gain and stop-loss non-reference alleles within each accession is provided in Supplementary Data 7. We observed a clear pattern in the genetic load reflecting differences between the Andean and Mesoamerican origins, with the Andean accessions showing a higher genetic load due to the bottleneck before domestication (Fig. 4d). We observed a reduced genetic load in EU_AND for both the loss-of-function (EU_AND *vs* AM_AND; Mann-Whitney p=0.03) and the missense mutations (EU_AND *vs* AM_AND; Mann-Whitney p=0.005). Conversely, we did not observe a reduction of genetic load in EU_MES for both missense and loss-of-function mutations (Fig. 4d). This suggests that the relatively short period of inter-genepool hybridisation, followed by selfing and recombination, promoted the purging of deleterious alleles accumulated in the European Andean pool. The role of hybridisation and subsequent recombination was also supported by the pattern of long-range LD in Europe compared to the Americas (Fig. 4c). The pattern for private alleles (i.e., those not identified in other genepools or populations; Supplementary Note 29) in the American and European accessions for low-frequency mutations (< 5%) revealed a 1.44-fold higher frequency of non-synonymous over synonymous mutations in Europe (Supplementary Fig. 61). This may have resulted from the pattern of crop dissemination, which was probably characterised by the exchange of small quantities of seeds and several sequential bottlenecks, followed by rapid population growth at the single-farm level, leading to the fixation of most mutations due to the small population size (i.e., a founder effect). In this demographic context, most mutations would be fixed rapidly at the local level (within the population grown by a single farmer). However, it is also possible that the purging of deleterious mutations, due to hybridisation following seed exchange among farmers and the co-occurrence of different varieties in the same fields^21^, facilitated the combined effects of natural and human selection against deleterious recessive alleles and the capture of valuable variants.

### Selection and adaptive introgression

We defined putative adaptive introgressed loci (PAIL) as those showing signatures of adaptive introgression meeting the following requirements: (a) an excess of introgression based on Chromopainter, (b) a signature of selection detected using the hapFLK method, which analyses multiple populations, jointly considering their hierarchical structure^37^, and (c) an outlier *F_ST_* value between Europe and the Americas, suggesting different patterns of diversity between these regions. The level of genetic differentiation (*F_ST_*) among genetic groups is reported in Supplementary Tab. 14 (see also Supplementary Fig. 62-63 and Supplementary Note 30 for details) The regions showing an excess of introgression (see Supplementary Tab. 15 and Supplementary Data 8) have been identified as described in Supplementary Note 31. Although the hapFLK method allowed us to identify selection signatures across the genome, an outlier *F_ST_* value was used to define which selection signature represents significant differentiation at the genomic level between American and European populations, suggesting selection in Europe. The identification of excess introgression independently of hapFLK provides evidence for adaptive introgression and the identification of PAIL. We also considered the occurrence of inter-chromosomal LD across genomic regions, private to European genepools (Fig. 4e, Fig. 5, see also Supplementary Note 32, Supplementary Fig. 64-66, Supplementary Data 9-12 and Supplementary Tab. 16) as an interesting signal to define regions potentially involved in adaptive processes. We also identified, through a GWA analyses (see Supplementary Notes 33-36)genomic regions associated to flowering and growth habit (Supplementary Fig. 67-75 and Supplementary Tab. 17). Finally, we suggest a putative role for candidate loci/ genes, combining our data on adaptive introgression regions and GWAS peaks with a literature search on the orthologous genes function (see Supplementary Notes 37-52 for details). Adaptive introgression appears particularly important for the evolution of the European genotypes of Mesoamerican origin (EU_MES). We identified 44 Andean genomic regions with excess introgression (23 of which showed signals of adaptive introgression) that are shared by > 90% of the European genotypes, spanning all chromosomes except PvChr11 (Supplementary Data 8; F(AND)) and ranging from ∼5 to ∼118.5 kb in length. An Andean allele frequency of 96% was detected along a genomic segment of PvChr01 (Chr01:46175616–46294040; Supplementary Data 8) that shows signs of adaptive introgression. This region contains 18 genes including *Phvul.001G203400*, which is orthologous to *OVARIAN TUMOR DOMAIN-CONTAINING DEUBIQUITINATING ENZYME 5* (*OTU5*) (see Supplementary Note 43 and Supplementary Data 13; row 16). In *Arabidopsis thaliana,* the function of this gene is to recalibrate and maintain cellular inorganic phosphate homeostasis^38–39^. The common bean orthologue may therefore be involved in the phosphate starvation response, making it an interesting candidate for further testing. In the same region on chromosome Pv01, we also identified *Phvul.001G204600* and *Phvul.001G204700* (see Supplementary Notes 41 and 50 and Supplementary Data 13; rows 29 and 30), which are orthologous to *LUMINIDEPENDENS* and *NUTCRACKER*, respectively (Fig. 5).

**Figure 5.**
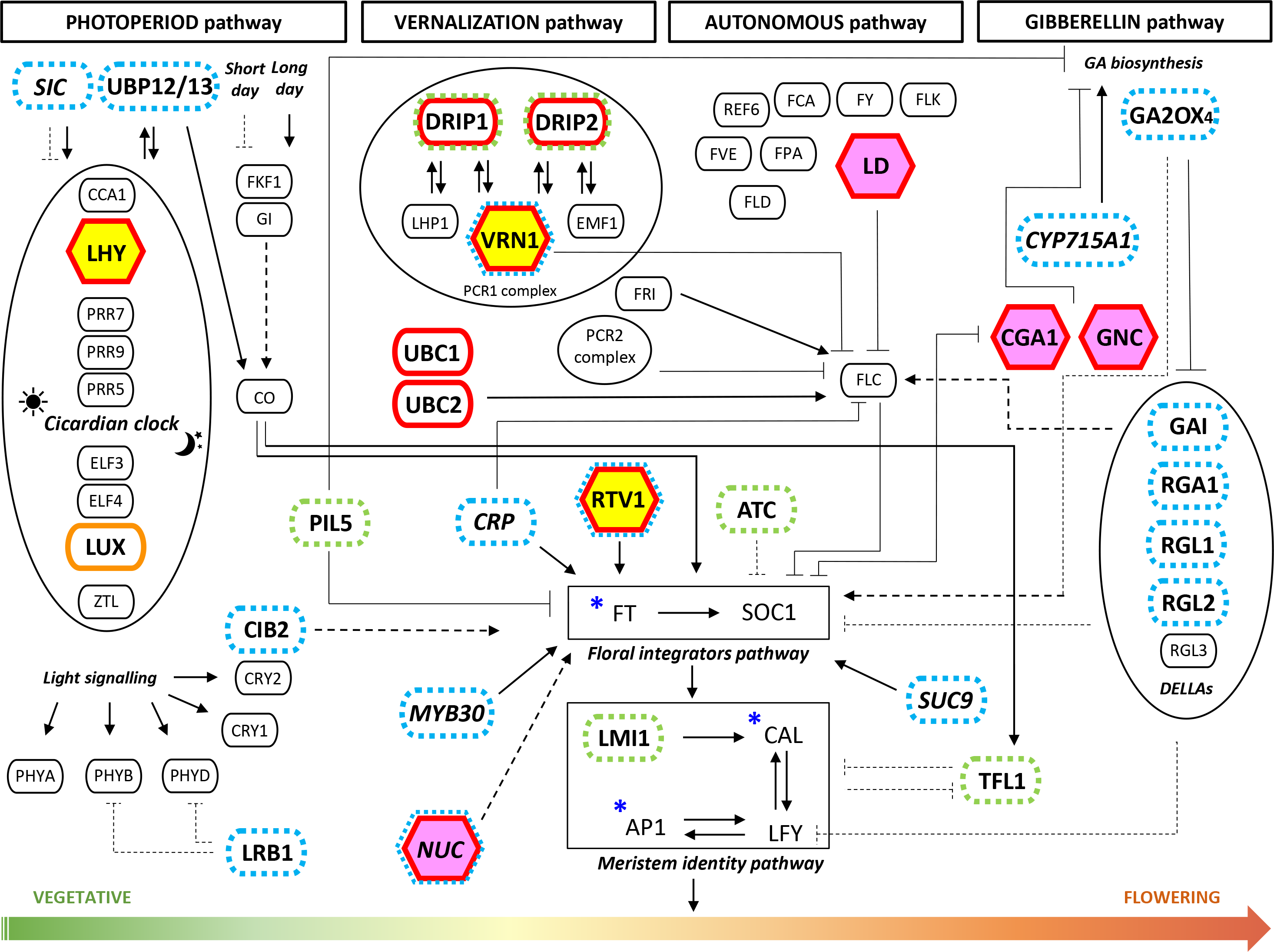
Candidate genes for adaptation. Schematic representation of the regulatory networks underlying the four major flowering pathways in *Arabidopsis thaliana*. The genes involved in the photoperiod, vernalisation, autonomous and gibberellin pathways that lead to the transition from vegetative to flowering are shown below the corresponding pathway. Additional genes belonging to secondary pathways that interact with the main regulatory flowering networks are shown in italic. Orthologues genes in common bean showing signatures of adaptive introgression, and those located in GWA peaks in our study are highlighted as follows: yellow hexagons – common bean orthologues of *LHY* (*Phvul.009G259400*, *Phvul.009G259650*) and VRN1 and *RTV1* (*Phvul.011G050600*) showing private inter-chromosomal linkage disequilibrium (LD) in the EU_M pool; pink hexagons – common bean orthologues of *LD* (*Phvul.001G204600*), *NUC* (*Phvul.001G204700*), *CGA1* and *GNC* (*Phvul.003G137100*) showing private inter-chromosomal LD in the EU_M pool ; red outlines – at least one orthologous gene in common bean showing signature of selection, introgression and with a significant differentiation (*F_ST_* index) between American and European accessions (p < 0.05); orange outlines – at least one orthologous gene in common bean showing a signature of selection with no significant *F_ST_* (p < 0.05); blue asterisks – at least one orthologous gene in common bean showing a signature of introgression; dashed blue outlines – at least one orthologous gene in common bean located within 50 kb centered on a significant GWA peak for days to flowering; dashed green outlines – at least one orthologous gene in common bean located within 50 kb centered on a significant GWA peak for growth habit; arrows – positive regulation of gene expression; truncated arrows – repression of gene expression; solid lines – direct interactions; dashed lines – indirect interactions in *A. thaliana*. Candidate genes for adaptation or post-domestication of the common bean in Europe, orthologous to those involved in flowering-related pathways, are shown in parentheses: *UBP12/13* (*Phvul.007G234000*); *LHY* (*Phvul.009G259400*, *Phvul.009G259650*); *LUX* (*Phvul.011G062100*); *PIL5* (*Phvul.001G168700*); *CIB2* (*Phvul.008G133600*); *LRB1* (*Phvul.006G109600*); *DRIP1/2* (*Phvul.001G157400*, *Phvul.007G177500*); *VRN1*, *RTV1* (*Phvul.011G050600*); *UBC1/2* (*Phvul.003G191900*); *LD* (*Phvul.001G204600*); *TFL1*, *ATC* (*Phvul.001G189200*); *GA2OX4* (*Phvul.006G120700*); *CGA1*, *GNC* (*Phvul.003G137100*); *GAI*, *RGA1*, *RGL1*, *RGL2* (*Phvul.001G230500*); *LMI1* (*Phvul.001G184800*, *Phvul.001G184900*); *SIC* (*Phvul.008G182500*); *CRP* (*Phvul.008G142400*); *MYB30* (*Phvul.008G041500*); *NUC* (*Phvul.001G154800*, *Phvul.001G204700*, *Phvul.011G074100*); *SUC9* (*Phvul.004G085100*, *Phvul.004G085400*, *Phvul.004G085594*); *CYP715A1* (*Phvul.007G071500*).

This adaptive introgression region in the common bean genome is close to known regions associated with flowering time, such as the *fin* locus controlling determinacy, and explaining phenotypic variation also for flowering time^40^. The region also co-maps with *Phvul.001G189200* (*PvTFL1y*; Pv01:44856139–44857862)^41^, and shows linkage^40^ to the *Ppd* locus controlling photoperiod sensitivity^42^. Wu *et al*.^25^ recently identified several markers on chromosome Pv01 associated with flowering under different conditions, with one at ∼45.5 Mb on Pv01. In our GWA, we identified a significant association between photoperiod, flowering time (see also Supplementary Note 36) and the marker S01_48049738, which is found ∼400 kb downstream of *Phvul.001G221100* (Chr01:47642033– 47647745) a gene that has been proposed as a candidate for the common bean *Ppd* locus^43, 44^.

Overall, we identified 77 genes that are PAIL. These represent ∼11% of all genes (n = 681) showing signatures of selection in Europe (i.e., selection signatures identified with hapFLK and being in an *F_ST_* outlier window; n = 354) and/or excessintrogression (n = 404). Accordingly, 277 genes show selection in Europe but not excess introgression, and 327 show excess introgression but not selection in Europe. The 77 PAIL show enrichment in seven Gene Ontology categories including GO:0048523, negative regulation of cellular processes; GO:0010228, vegetative to reproductive phase transition of the meristem; GO:0042445, hormone metabolic processes; GO:0009657, plastid organisation; GO:0042440, pigment metabolic processes; GO:0009733, response to auxin; and GO:0070647, protein modification by small protein conjugation or removal (see Supplementary Note 39 and Supplementary Fig 76). Enrichment analysis also suggested that flowering has been an important target of adaptive introgression, highlighting the importance of genes that may be associated with adaptation to abiotic and biotic stress.

Adaptive introgression signals were identified in many *P. vulgaris* genes with a putative role related to flowering (Supplementary Notes 40 and 41), including orthologues of genes involved in the four major *A. thaliana* flowering pathways (Fig. 5). Significant examples include *Phvul.009G259400* and *Phvul*.009G259650 (see Supplementary Note 41 and Supplementary Data 13; rows 90 and 92), which are orthologues of *A. thaliana LATE ELONGATED HYPOCOTYL* (*LHY*), both located within the same adaptive introgressed region of chromosome PvChr09 (characterised by an Andean allele frequency of 96% in the European genotypes; Supplementary Data 8). Notably, the transcription factor encoded by *LHY* is a pivotal oscillator in the morning stage of the circadian clock. It is connected to the indirect suppression of the middle, evening, and night complex genes by *CIRCADIAN CLOCK ASSOCIATED 1* (*CCA1*)^45^ (Fig. 5), with a putative function in regulating flowering that has also been proposed in the common bean^46^. In the EU_MES population, these two *LHY* orthologues show private and significant inter-chromosomal LD with *Phvul.011G050600*, which yields GWA signals for flowering time and adaptive introgression (see Supplementary Note 50 and Supplementary Data 13; row 97). The latter is an orthologue of the *A. thaliana* genes *VERNALISATION 1* (*VRN1*) and *RELATED TO VERNALISATION1 1* (*RTV1*) (Fig. 5), which are needed to activate the floral integrator genes following long-term exposure to cold temperatures^47^. The inter-chromosomal LD between these putative flowering genes, which is private to the EU_MES accessions, may be the result of epistatic selection. Analogous examples include *Phvul.001G204600* and *Phvul.001G204700* (see Supplementary Notes 41 and 50 and Supplementary Data 13; row 29 and 30), which are orthologous to *LUMINIDEPENDENS* (*LD*) and *NUTCRACKER* (*NUC*), respectively. Both are located in a region of PvChr01 as described above, and are in private inter-chromosomal LD with *Phvul.003G137100* (Supplementary Data 13; row 38) on PvChr03, which is orthologous to *GATA, NITRATE-INDUCIBLE, CARBONMETABOLISM INVOLVED* (*GNC*), and *CYTOKININ-RESPONSIVE GATA FACTOR 1* (*CGA1*). *LD* is one of the eight genes identified so far in the *A. thaliana* autonomous pathway and it represses *FLOWERING LOCUS C* (*FLC*) to promote the transition from vegetative growth to flowering (Fig 5). *NUC* encodes a transcription factor that positively regulates photoperiodic flowering by modulating sugar transport and metabolism via the *FLOWERING LOCUS T* (*FT*) gene^48, 49^. The paralogous *GNC* and *CGA1* genes act redundantly to promote greening downstream of the gibberellin signalling network^50^.

## Discussion

We have shown that adaptive introgression was necessary for the successful dissemination and adaptation of the common bean in Europe. We combined genome resequencing, metabolomics, classical phenotyping and data analysis for chromosome-level genetic assignment and environmental association. Our data indicate that the Andean genepool was the first to be successfully introduced in Europe, most likely from Francisco Pizarro’s expedition to northern Peru in 1529. Most of the Andean genetic background of the European common bean was contributed by the A1/Nueva Granada and A3/Chile races after secondary domestication, whereas the more photoperiod-sensitive A2/Peru race contributed little to European common bean germplasm. The secondary domestication of these Andean races, related to the latitudinal expansion of the cultivation areas from the Andean centres of origin, facilitated the successful dissemination of the Andean common bean in the Old World. However, we propose that the adaptive introgression observed in Europe for individuals mainly belonging to the M1/Durango-Jalisco race was an important event that underpinned the successful dissemination of this Mesoamerican ancestry in Europe. Indeed, genomic analysis indicated that Andean genotypes were rapidly disseminated, whereas Mesoamerican genotypes were eventually disseminated in Europe following introgression from the Andean types. In addition to the flowering time data gathered from the European and American accessions, we also identified clear signatures of selection in common bean orthologues of genes representing the major flowering pathway and environmental adaptations, such as the *OTU5* gene involved in the inorganic phosphate starvation response. These are interesting candidate loci for further validation. Finally, we propose that the dissemination of common bean accessions in Europe may have been influenced by political factors and constraints in the 16^th^ century, including the interaction between political and religious powers in Western Europe and the subdivision of the European continent into Islamic and Christian countries.

## Methods

### Plant materials

Original seeds for 218 common bean accessions (*P. vulgaris*) were collected from international gene banks or individual institutional collections. We produced 199 single seed descent (SSD) lines by performing at least three cycles of self-fertilisation. For the remaining 19 accessions, one seed per accession was sampled directly from original seeds provided by the donor.

### Experimental design and phenotyping

Plants were grown across 10 different environments in fields and greenhouses, applying long-day (7), short-day (2) and intermediate photoperiod conditions. During the summers of 2016 and 2017, four field trials were carried out in Italy (Villa d’Agri, Marsicovetere, Potenza) and in Germany (Gatersleben IPK). Six additional greenhouse experiments were performed under controlled conditions in Golm (Potsdam, Germany), Potenza (Italy) and Villaviciosa (Spain) during 2016, 2017 and 2018 (Supplementary Notes 3 and 4).

Classical phenotyping was carried out on the 199 SSD lines, focusing on two main traits: days to flowering (DTF), defined as the number of days from sowing until 50% of plants showed at least one open flower; and growth habit (GH), defined as determinacy *vs* indeterminacy on a single plant basis. Photoperiod sensitivity (PS) was calculated as the ratio between DTF in long-day and short-day experiments.

Descriptive statistics were calculated for the different phenotypic traits using R (https://cran.r-project.org/) or JMP 7.0.0^51^. The restricted maximum likelihood (REML) model implemented in JMP 7.00 was used to calculate least square means (LSM) and the best linear unbiased predictors (BLUPs) of each genotype. The REML model was also used to calculate the broad-sense heritability (h^2^B) for each quantitative trait by considering genotypes and environments as random effects. The distribution of DTF in each environment and Pearson’s pairwise correlation between environments were calculated using the *corrplot* and *PerformanceAnalytics* R packages^52, 53^.

Molecular phenotyping (see Supplementary Note 5) of the 199 accessions was performed on the first trifoliate fully expanded leaves harvested under long-day conditions in three biological replicates. Secondary metabolites were measured as described in Perez de Souza *et al*.^54^. For non-targeted metabolomics, chromatogram peak detection and integration were achieved using GeneData REFINER MS 10.0 (http://www.genedata.com). To explore the molecular phenotypic diversity, we performed non-targeted metabolic fingerprinting by high-throughput LC-MS analysis. Mass signals that were not detected in ≥ 50% of the samples and/or those with a peak intensity ≤ 1000 were excluded. Heritability was analysed as stated above, setting genotype and continent as random effects. The heritability was calculated based on 190 accessions (94 from the Americas and 96 from Europe) with more than one replicate.

### Sequencing, variant calling and annotation

Genomic DNA was extracted from frozen young leaves of the 199 SSD lines grown in greenhouse and directly from seeds of the remaining 19 accessions using the DNeasy Plant Mini Kit (Qiagen, Hilden, Germany). It was sheared using a Covaris E220 device to fragments of ∼550 bp and PCR-free libraries were constructed using the KAPA HyperPrep Kit. Paired-end sequencing libraries were sequenced on Illumina HiSeq2500 or HiSeq4000 devices and labelled with different barcodes.

Sequencing data were aligned to the common bean reference genome v2.0^15^ using BWA-mem v0.7.15^55^. Unmapped reads were mapped to the *P. vulgaris* chloroplast genome (NCBI NC_009259). In both cases, SNPs were called using SAMtools^56^ and Genome Analysis ToolKit (GATK) v3.6^57, 58^. In SAMtools, duplicated reads were removed with *rmdup* and SNPs were discovered with *mpileup* for filtered high-quality alignments (–q 10) and bases (–Q 20) and then genotyped with BCFtools^59^. In GATK, duplicated reads were sorted and filtered with Picard v2.4.1 (http://broadinstitute.github.io/picard). Variants were then called according to GATK best practices and pre-filtered using the recommended parameters for hard filtering. Chromosomal and overlapping SNPs reported by both methods were retained and the genotypes produced by GATK were selected. We applied additional filtering with VCFtools^60^ (--minDP 3 --max-missing 0.5 --maf 0.05) and excluded SNPs with proportions of heterozygous genotypes > 0.01. SNPs were annotated with SnpEff v4.3s^61^.

### Population structure analysis

Population structure analysis was applied to the SAMtools/GATK overlapping SNP callset, followed by filtering to retain only genomic positions with a QUAL score ≥ 30 and a global depth of coverage between 1/3 and 4 times the mean value. Individual genotypes called using two reads or fewer were marked as missing data. Imputation and phasing were performed with Beagle^62^. ADMIXTURE v1.3^13^ was used for population structure analysis. The unphased variants were filtered by taking one SNP every 250 kb using VCFtools. In ADMIXTURE, we varied K from 1 to 20 in 20 replicates and applied the analysis independently over the whole sample of American and European (n = 218) accessions or using the American accessions only (AM, n = 104). We dealt with potential cryptic population structures within each pool as previously described^16, 31, 32^.

Population structure was inferred from chloroplast data using Bayesian Analysis of Population Structure (BAPS) v5.3^63, 64^. Mixture analysis was used to determine the most probable K value according to the data. *Clustering with linked loci* analysis was chosen to account for the linkage between sites. Ten repetitions of the algorithm for each K value from 2 to 20 were applied. The relationships between genotypes were determined using the neighbour-joining method in MEGA X^65^ with a bootstrap value of 1000. Gaps and missing data were excluded.

### Chromosome painting

Chromosome painting was applied to the phased variants in ChromoPainter v2.0^20^. The effective population size (Ne) and mutation rates (Mu) were estimated individually for each accession using 10 iterations of the expectation–maximisation algorithm in ChromoPainter. The estimated parameters were fixed in a new round of analysis producing the final chromosome painting of the recipient haplotypes. Donor individuals were chosen as follows, according to their ancestry proportion inferred by admixture: (a) Mesoamerican individuals with a q value > 0.99 in the admixture run (K = 3 using all American accessions), and (b) Andean individuals with a q value consistently > 0.99 from K = 2 to K = 4 in the admixture run restricted to Andean accessions. Donors were subdivided into the five groups inferred by ADMIXTURE (AM_M1, AM_M2, AM_A1, AM_A2 and AM_A3) and were used to estimate their contribution to the ancestry of each SNP in the recipient individuals. Individual SNP probabilities were then combined in 10-kb non-overlapping sliding windows along chromosomes and each window in each recipient haplotype was assigned to one of the five donor groups if a probability ≥ 0.8 was observed (see also Supplementary Note 19). The total proportion of genetic material from the seven groups or “unknown” (genotypes assigned to none of the groups) was computed for each recipient individual and for each chromosome (both pairs). The final assignment of each recipient accession to the genepools was based on (a) the total proportion of windows attributed to Mesoamerica or Andes, and (b) the number of chromosomes assigned to the two genepools following the majority rule criterion.

The attribution of each genomic window to the seven groups was also used to estimate the length of the introgressed blocks within each European accession. Each haplotype of the EU_AND accessions was traversed, merging consecutive windows attributed to any of the Mesoamerican clusters. Bedtools^66^ was used to join windows within a maximum distance between elements of 50 kb to deal with artificially broken introgressed blocks. The length of each Mesoamerican block in each EU_AND individual was recorded for each chromosome and was then filtered to remove blocks composed of single windows (10 kb). The final within-individual distribution of lengths was characterised by the median value due to the non-normal distribution of the data.

For spatial analyses, the ecological data (resolution ∼1 km^2^) were downloaded from WorldClim (http://www.worldclim.org)67 for a total of 19 bioclimatic variables and 24 monthly variables (Supplementary Data 6). The vegan *R* package^68^ was used to calculate the geographical and ecological distances, the Mantel statistics, and the spatial autocorrelation. Initially, the Mantel statistics were tested by 10^3^ permutations and the autocorrelogram was calculated between 10 distance classes of nearly 540 km each, determining the significance of the correlation in each class by 9999 permutations. We then applied environmental association analysis with a multivariate correlation analysis between the p values (proportion of membership to the five genetic groups) assigned to each European accession and the ecological variables registered at the collection site.

### Genetic diversity

The genetic diversity within groups of accessions, defined according to their geographic origin and genepool, was quantified using the theta estimator θπ^69^. The --site-pi VCFtools flag was used to obtain a per-SNP estimate that was subsequently filtered according to the genome annotation, including only positions located: (a) in callable regions, (b) in coding regions, and/or (c) in neutral regions (Supplementary Note 24). The per-site θπ estimate was then summed up and divided by the size of each specific region to calculate a global estimate. A raw estimate of θπ along chromosomes, averaged over 100-kb non-overlapping windows, was also computed to highlight chromosomal regions with different levels of genetic diversity. To evaluate the stability of the θπ estimate at different missing data levels, a masked dataset was obtained by filtering introgressed alleles (identified by ChromoPainter) and alleles with an ambiguous assignment, within European accessions. The --site-pi and --missing-site commands in VCFtools were used to obtain a per-site θπ estimate and the proportion of missing data for each position, respectively. The global within-group θπ was computed for the callable, coding and neutral genomic partitions, excluding regions with an average (over SNP) minimum mean proportion of non-masked individuals (PIND) from 0% to 100%.

To detect patterns of private alleles, missense and synonymous variants were screened in American and European accessions (Supplementary Notes 28 and 29). Variants that were private to the European or American groups were retained and divided into those with low (< 5%) and medium-high (> 5%) within-sample frequencies. The genomic coordinates related to private alleles segregating at different frequencies in the American and European groups of accessions were intersected with the gene annotations, and the burden of missense and synonymous mutations was recorded for each gene element.

The magnitude of the genomic differentiation between and within America and Europe was evaluated using the Weir & Cockerham estimator *F* ^70^. We estimated the baseline differentiation between and within the two continents. In addition, the *F_ST_* was then computed in 10-kb non-overlapping sliding windows between each pair of groups using VCFtools. The mean and the interquartile range (IQR) of the windows-based distribution were used as a point estimate of the differentiation between groups and to evaluate its dispersion.

### Comparison of the genetic structure, molecular phenotype and flowering data

Analysis of variance (ANOVA) was used to evaluate differences between the genetic subgroups using the first principal component related to DTF and PS (see also Supplementary Notes 6 and 23) as representative phenotypic traits. Principal component 1 was obtained from the secondary metabolites with a high hereditability of H^2^ > 0.65 (Supplementary Notes 5 and 23) and this was used as a phenotype for comparison between the genetic subgroups.

### Tagging the signatures of adaptation in Europe

The presence of excess of introgression and selection was investigated in Europe (Supplementary Notes 21 and 31). To detect deviations from the frequencies expected in the absence of demographic and selection forces, the ChromoPainter output was parsed by tracing the assignment of each SNP to the corresponding Mesoamerican or Andean groups. For each SNP, we computed the proportion of haplotypes assigned to the Mesoamerican or Andean groups. We extracted the genomic coordinates of SNPs showing an unexpected proportion of Andean alleles (threshold: EU_A, 71 × 2 = 142 haplotypes, plus 50% of the Mesoamerican ones, EU_M, 43 × 2 × 0.5 = 43, Fobs ≥ 0.811). The putative SNP targets of Mesoamerican introgression events were identified according to the same rationale (threshold: EU_M, 43 × 2 = 86, plus the 50% of Andean ones, EU_A, 71 × 2 × 0.5 = 71, Fobs = 0.688). The Bedtools slop -b 2500 and merge -d 10000 functions were used to pass from SNP point coordinates to 5-kb regions and then merge into larger genomic blocks if the relative distance between them was < 10 kb. Only genomic regions supported by at least three SNPs were retained.

The hapFLK^37^ method was used to identify selection signatures. The local genomic differentiation along chromosomes, as measured by haplotypic *F_ST_*, was compared to the expectation given by the inferred genomic relationships between groups, considering the genetic drift within groups. Accessions were subdivided into the AM_A (n = 30), AM_M (n = 36), EU_A (n = 71) and EU_M (n = 43) groups and VCFtools was used to sample a single SNP every 250 kb. This set of SNPs was used to construct a neighbour-joining tree and a kinship matrix according to the Reynolds’ genetic distance matrix between the four groups of accessions, constituting a genome-wide estimate of population structure. The hapFLK statistics were then computed independently on each chromosome over the complete SNP dataset and averaged over 20 expectation–maximisation cycles to fit the LD model. Initial analysis fixed the number of haplotype clusters to five. A second run was conducted, selecting the appropriate number of haplotype clusters based on the fastPHASE^71^ cross-validation procedure, implemented in the *imputeqc* R package (https://github.com/inzilico/imputeqc). VCFtools was used to extract a subset of SNPs spaced at least 100 kb apart on each chromosome, and five independent copies of the SNP set were generated, randomly masking 10% of the variants. We then used fastPHASE v1.4.8 to impute the missing genotypes in each dataset, setting the number of haplotype clusters to 5, 10, 20, 30, 40 and 50. The EstimateQuality function was used to compute the proportion of wrongly imputed genotypes (Wp) for each combination, and the K value, minimising the mean Wp proportion across the five SNP set replicates, was selected as the most supported number of haplotype clusters. The analysis was replicated using all or only American accessions. The scaling_chi2_hapflk.py script was used to scale hapFLK values and compute the corresponding p values. The significant SNPs (p < 10^-3^; FDR < 0.05) were extracted and bedtools was used to create a 10-kb region centred on each significant SNP and to merge overlapping regions within a maximum distance of 5 kb. The two sets of regions were merged forming the extended set (constituted by the union of the two sets) and the restricted set (containing only regions supported by both runs). To pinpoint putative regions under selection in Europe, the extended and restricted sets were intersected with the *F_ST_* window analysis, and only regions containing at least one *F_ST_* window located in the top 5% or top 1% were retained.

### Linkage disequilibrium (LD)

The relationship between LD and physical distance along chromosomes was evaluated in America and Europe, and successively within the American subgroups. PopLDdecay^72^ was used to compute the correlation (r^2^) between allele frequencies at pairs of SNPs along the chromosomes, setting a minimum minor allele frequency of 0.1 and a maximum distance between SNPs of 5 Mb.

The level of inter-chromosomal LD was also evaluated. VCFtools was used to sample one SNP every 10 kb and compute r^2^ between pairs of markers located on different chromosomes. The analysis was performed independently over the American subgroups, using only SNPs that were segregating within each group of accessions with a minor allele frequency > 0.05, and only pairs of SNPs showing an r^2^ value ≥ 0.8 were retained. Multiple pairs of SNPs pointing to the same chromosomal regions were merged if within a distance of 100 kb from each other and only pairs of regions spanning at least 500 kb on each side were retained. The whole analysis was also repeated including only SNPs falling in the putative regions under selection, decreasing the minimum width of retained regions from 500 kb to 50 kb. Link plots showing high-LD regions were produced using Rcircos^73^.

### Genome-wide association study (GWAS)

GWAS was carried out for the growth habit, flowering time and photoperiod sensitivity data (see Supplementary Notes 6, 33-36). First, we ran a single-locus mixed linear model (MLM) in the R package MVP^74^ (https://github.com/XiaoleiLiuBio/MVP). Bonferroni correction at α = 0.01 was applied as the significance threshold for each trait. The analysis was then conducted using the multi-locus stepwise linear mixed-model (MLMM)^75^ (https://github.com/Gregor-Mendel-Institute/MultLocMixMod). By applying a stepwise approach, this includes the most significant SNPs as cofactors in the mixed model. The mBonf criterion was used to identify the optimal results with Bonferroni correction at α = 0.05.

### Investigation on the putative function of candidate genes for adaptation

Common bean genes orthologous to *A. thaliana* and legume genes were identified using Orthofinder^76, 77^ (Supplementary Note 37). The putative function of poorly characterised genes was predicted based on orthologous relationships and literature screening. The orthologue and known genes involved in DTF, GH and PS were screened against the GWAS results. Genes within 50 kb on either side of each significant SNP associated with DTF and GH, and genes located within selection scan and introgression scan regions, were investigated by GO term enrichment analysis (biological process, cellular component and molecular function) using Metascape^78^ (http://metascape.org).

### Data availability

The raw sequence reads generated and analysed in this study have been deposited in the Sequence Read Archive (SRA) of the National Center of Biotechnology Information (NCBI) under BioProject number PRJNA573595. Source data are provided with this paper as Source Data file. DOIs for the BEAN_ADAPT PV-core 2 accessions are provided in the Supplementary Data 1. Data from the databases/ data repositories Phytozome v.2.1, NCBI NC_009259 (*P. vulgaris* chloroplast genome), WorldClim (http://www.worldclim.org), https://zenodo.org/record/3236786#.Y49hDXbMK3A (vcfs for the whole genomes sequences data from Wu *et al*.^25^) have been used and analyzed within this manuscript.

## Code availability

Codes are available in the https://github.com/PapaLab/Bean_Adapt repository.

## Acknowledgements

This manuscript is dedicated to our former collaborator Dr. Monica Rossi, who passed away at the age of 44 in 2019, and to Dr. Chris Berry, a colleague and close friend who contributed to the editing of an earlier version of this manuscript and who recently passed away. This work was carried out as part of the BEAN ADAPT project, funded by the ERA-CAPS Programme (2014 Call, Expanding the European Research Area in Molecular Plant Sciences). E.M., V.D. and E.B. acknowledge funding from the European Union’s Horizon 2020 research and innovation programme under grant agreement No. 774244 (BRESOV Project), G.C. and G.F. acknowledge funding from the European Union’s Horizon 2020 research and innovation programme under grant agreement No. 862862 (INCREASE Project), S.A. and A.R.F acknowledge funding from the European Commission for the PlantaSyst project (SGA-CSA nos. 664621/739582 under FPA no. 664620).

## Author Contribution

A.G., P.G., A.R.F., S.A.J. and R.P. conceived and managed the project; E.Be. and R.P. wrote the article; E.Be., A.B., C.X., E.Bi., M.R., S.A., V.D.V., K.N., G.C., G.F., P.L.M., A.G., P.G., A.R.F., S.A.J. and R.P. contributed to the writing and drafting; E.Be., A.B., C.X., E.Bi., M.R., S.A., V.D.V., T.G., K.N., G.C., G.F., L.N., J.J.F., A.C., G.A., P.L.M., G.B., A.G., P.G., A.R.F., S.A.J. and R.P. contributed to the editing of the article; E.Be., A.B., C.X., E.Bi., M.R., S.A., V.D.V., T.G., K.N., G.C., G.F., E.M., E.T., L.N., A.A., C.L., J.J.F., A.C., P.L.M., GB, A.G., A.R.F., S.A.J. and R.P. contributed to the writing and organisation of the supplementary material; E.Be., E.Bi., S.A., T.G., K.N., L.N., G.L., L.J., J.J.F. and A.C. performed DNA extraction, field and greenhouse experiments; C.X., J.H.S. and S.A.J. conducted sequencing and primary bioinformatic analysis; S.A. and A.R.F. conducted metabolomics analysis; A.B., E.Bi., M.R., G.F., E.T., C.L., G.B. and R.P. contributed to data analysis; E.Be., A.B., C.X., E.Bi., M.R., S.A., V.D.V., G.B., A.R.F., S.A.J. and R.P. contributed to coordinate data analysis and data integration. All authors read and approved the article.

## Competing Interests

The authors declare no competing interests.

## Figure legends

## Notes

### Competing Interest Statement

The authors have declared no competing interest.

### Summary of Updates

The manuscript is going through a final revision process before being published on a peer review Journal, and we would like to keep it updated also on this platform

